# Long-read based assembly and annotation of a *Drosophila simulans* genome

**DOI:** 10.1101/425710

**Authors:** Pierre Nouhaud

**Affiliations:** Institut für Populationsgenetik, Vetmeduni, Vienna, Austria

**Keywords:** genome assembly, PacBio, SMRT sequencing, IsoSeq

## Abstract

Long-read sequencing technologies enable high-quality, contiguous genome assemblies. Here we used SMRT sequencing to assemble the genome of a *Drosophila simulans* strain originating from Madagascar, the ancestral range of the species. We generated 8 Gb of raw data (~50× coverage) with a mean read length of 6,410 bp, a NR50 of 9,125 bp and the longest subread at 49 kb. We benchmarked six different assemblers and merged the best two assemblies from Canu and Falcon. Our final assembly was 127.41 Mb with a N50 of 5.38 Mb and 305 contigs. We anchored more than 4 Mb of novel sequence to the major chromosome arms, and significantly improved the assembly of peri-centromeric and telomeric regions. Finally, we performed full-length transcript sequencing and used this data in conjunction with short-read RNAseq data to annotate 13,422 genes in the genome, improving the annotation in regions with complex, nested gene structures.

## Introduction

DNA sequencing has experienced three revolutions which have profoundly impacted life sciences (Shendure et al. 2017). Forty years ago, Sanger et al. (1977) pioneered the development of a method which would allow to sequence genomes for the first time, starting with phages and culminating with humans. The cost and labor-intensiveness of this method led to the development of second-generation sequencing (SGS) technologies in the early 2000s. While Sanger sequencing produced reads of 600-1000 bp, SGS platforms would generate massive amounts of small reads (e.g., 125 bp for an Illumina HiSeq 2500) at a fraction of the price. SGS technologies are well-suited for resequencing studies but are limited for *de novo* genome assembly, since the length of DNA repeats is usually more important than the length of a single read. This issue would lead to assembly errors and fragmented genome assemblies (Alkan et al. 2011). The last revolution to date came from long-read, third-generation sequencing (TGS) technologies, which routinely produce reads of more than 10 kb (van Dijk et al. 2018). TGS can finally overcome the limitations associated with short reads for *de novo* genome assembly. However, their high error rate (> 10% for PacBio’s SMRT sequencing) requires a sequence correction step either by using SGS data, or by increasing TGS coverage. Another potential application of TGS with SMRT technology is full-length transcript sequencing. Transcript reconstruction from short reads often misses terminal exons or splice junctions (Steijger et al. 2013) and SMRT sequencing would provide better evidence for alternative splicing while improving the characterization of gene models (van Dijk et al. 2018).

The fruit fly *Drosophila simulans* has diverged from the model organism *D. melanogaster* 2-8 million years ago (Obbard et al. 2012). The first published *D. simulans* genome was sequenced at low coverage and was obtained from a mixture of strains (Begun et al. 2007; available on FlyBase: flybase.org). One of these strains (strain w501, from North America) was also sequenced at a deeper coverage (Hu et al. 2013). Palmieri et al. (2015) assembled the genome of a strain from Madagascar (strain M252), which represents *D. simulans* ancestral range (Kopp et al. 2006). All three assemblies were based on SGS and were comparable in terms of completeness.

Here, we report the sequencing and assembly of the *D. simulans* M252 genome using PacBio SMRT technology. We have also annotated the genome using ab initio predictions, RNAseq and full-length transcript sequencing using PacBio SMRT technology. These high-quality assembly and annotation will represent an important resource for population genomic studies in *D. simulans* and will help our understanding of genome evolution in *Drosophila*.

## Methods

### DNA extraction & long-read sequencing

We sequenced the *D. simulans* M252 strain provided by D.J. Begun and collected in 1998 by W. Ballard in Madagascar. This strain has been maintained in the lab through full-sib mating for several years and residual heterozygosity is expected to be low. High molecular weight DNA was extracted from a pool of larvae using a DNeasy Blood and Tissue Kit (Qiagen, Valencia, CA). DNA was sheared using a Covaris (Woburn, MA) g-TUBE and a single SMRTbell library was prepared using a DNA Template Prep Kit 1.0 and a DNA/Polymerase Binding Kit P6 v2 (Pacific Biosciences, Menlo Park, CA). This library was sequenced using 14 SMRTcells on PacBio RS II with C4 sequencing chemistry. Adaptors were removed and subreads filtered with SMRTanalysis version 1.4 using default parameters.

### De novo assembly

Third generation sequencing platforms have high error rates (> 10%, Rhoads & Au 2015) and a correction step is usually needed to build consensus sequences from raw reads. Most of the assemblers developed for these platforms implement this correction step, which requires substantial long read data coverage (at least 30x, Koren et al. 2017). As our sequencing depth was ~ 50× (see Results), we generated assemblies from subreads using different algorithms developed for long read data: HGAP in SMRTanalysis 1.4 (Chin et al. 2013), Falcon 0.3 (Chin et al. 2016), Canu 1.4 (Koren et al. 2017), miniasm 0.2 (Li 2016, no consensus step) and miniasm with Racon 0.5 for error correction (Vaser et al. 2017). Additionally we ran two hybrid assemblers, Spades 3.10 (Bankevich et al. 2012) and MaSuRCA 3.2.1 (Zimin et al. 2017) using long read data in conjunction with > 250× Illumina data previously generated from the same strain by Palmieri et al. (2015, 200 million 100bp paired-end reads sequenced on Illumina HiSeq2000). Apart from Falcon, which by default produced < 60 Mb assemblies (see File S1 for the final, optimized parameter file), all assemblers were run with default parameters, assuming when required an estimated genome size of 162 Mb (Bosco et al. 2007).

For each assembly, contigs were aligned to the *D. melanogaster* reference genome release 6.03 using the progressiveMauve algorithm from Mauve 2.4.0 (Darling et al. 2010). Since genome of both species are colinear (apart from a major chromosomal inversion in the 3R chromosome arm of the *D. melanogaster* iso-1 reference strain, which was reverse-complemented prior to this analysis), putative errors were detected when contigs aligned on two different *D. melanogaster* chromosome arms. Note that over all assemblies, no contig spanned centromeric regions.

We then selected the two best assemblies based on their total length, contiguity and absence of assembly errors (see Results). For each of them, two rounds of error correction were performed with Quiver in SMRTanalysis 1.4 using raw PacBio reads, and two additional rounds were done with Pilon 1.21 (Walker et al. 2014) using the aforementioned short read data from Palmieri et al. (2015). These two polished assemblies were combined using Quickmerge 0.2 (Chakraborty et al. 2016), and the same error correction pipeline was applied to the merged assembly.

Contigs from the merged assembly were anchored on the *D. melanogaster* genome r6 to create chromosome-level scaffolds using the nucmer module from the MUMmer 3.23 package (Delcher et al. 2002; Kurtz et al. 2004). Contigs were arranged into scaffolds using the show-tiling module from the MUMmer package following Nolte et al. (2013). The resulting assembly is hereafter referred to as *D. simulans* M252 genome version 2 (v2).

### Quality assessment

Assembly quality was assessed using an independent *D. simulans* sample derived from a pool of 202 isofemale lines collected in Florida and containing 200 million reads (population AP1 from Nouhaud et al. 2016, sequenced on Illumina HiSeq2000 with 100-bp paired-end reads, mean insert size: 374 bp). Reads were trimmed using the MottQualityTrimmer in ReadTools (Gómez-Sánchez & Schlötterer 2018, minimum quality: 20, minimum read length: 50 bp). Mapping was done successively against the *D. simulans* M252 genome version 1.1 (Palmieri et al. 2015), and against the version 2, using Bowtie2 2.2.6 (Langmead & Salzberg 2012, parameters ‐‐phred33 ‐‐end-to-end - X 1500) with DistMap (Pandey & Schlötterer 2013). For each resulting BAM file, we recovered the average percentage of nonproper pairs for non-overlapping, 10 kb sliding windows using the broken-pairs.pl script from Nolte et al. (2013). Nonproper pairs were defined as pairs were one of the mates (i) did not map, (ii) mapped to a different chromosome, (iii) mapped to the same strand as the other mate, or (iv) when distance between mates was greater than expected.

Genome completeness was assessed through BUSCO gene set analysis 3.0.2 (Simão et al. 2015; Waterhouse et al. 2018) using the Diptera gene set (2,799 orthologs).

### RNA extraction, long-read sequencing & IsoSeq pipeline

Flies from the M252 strain were maintained under a fluctuating thermal regime (12 hours under light at 28 °C, 12 hours under dark at 18°C) with density control (400 eggs per 300 ml bottle, containing 70 ml of standard *Drosophila* medium). After three generations, 50 males were frozen in liquid nitrogen in the middle of each temperature window. High quality RNA was extracted from each pool of male whole bodies using a RNeasy Plus Universal Mini Kit (Qiagen, Valencia, CA) and one SMRTbell library was built per sample (collected at 18 and 28°C, hence two libraries in total) following the Iso-Seq template preparation procedure from PacBio. Each library was sequenced using a single SMRTcell on PacBio Sequel with V2 chemistry at the VBCF NGS Unit (www.vbcf.ac.at). Raw data was processed using the IsoSeq pipeline (Gordon et al. 2015, available at https://github.com/PacificBiosciences/IsoSeq_SA3nUP) following PacBio’s guidelines. Briefly, the different steps are:

1. Identification of full-length (FL) reads for which all 5′-primer, polyA tail and 3′-primer have been sequenced;
2. Clustering of FL reads at the isoform level;
3. Alignment of non-FL reads to the isoform clusters;
4. Error correction using both FL and non-FL reads with the Arrow model.

This de novo pipeline outputs FASTQ files containing two sets of error-corrected, full-length isoforms: the high-quality set contains isoforms supported by at least 2 FL reads with an accuracy of at least 99%, while isoforms from the low-quality set display an accuracy < 99% (reasons could be insufficient coverage or rare transcripts). The pipeline was run first by pooling both libraries (giving a high-confidence set of transcripts for annotation, see below) and then for each library independently.

### Annotation

Repeats in the v2 genome were masked with RepeatMasker (http://www.repeatmasker.org/) after initial training based on *D. melanogaster.* We then used Maker 2.31 (Cantarel et al. 2008) to annotate the genome by first incorporating in silico gene models detected by Augustus 3.3 (Stanke & Morgenstern 2005), a de novo *D. simulans* transcriptome built with RNA-Seq data from Palmieri et al. (2015, single library combining multiple developmental stages and sequenced on Illumina HiSeq2000 with 100-bp paired-end reads, see initial publication for details) using Trinity r2014-07-17 (Haas et al. 2013) and all RefSeq protein sequences available for *D. simulans* on NCBI (n = 295,428, accessed on 31/10/2017), along with protein sequences from *D. melanogaster* r6.15 obtained from FlyBase (Gramates et al. 2017). The software was allowed to take extra steps to identify alternate splice variants.

Visual inspection of the output and comparison to the *D. melanogaster* annotation showed many gene fusion events and incorrect reconstruction of nested gene structures (see Results and Fig. 2). No significant improvement was detected after (i) decreasing the physical distance used to extend evidence clusters sent to gene predictors (pred_flank parameter, set to 100 bp instead of default 200 bp) and (ii) limiting the use of RNA-Seq data during the annotation to reduce gene fusion events (correct_est_fusion parameter disabled), as recommended in the Maker documentation.

To solve this issue, we included the IsoSeq data in the annotation procedure. The IsoSeq pipeline was ran simultaneously on the two combined libraries. The two resulting FASTQ files containing low and high quality sets of isoforms were pooled and aligned against the v2 genome using gmap r2017-10-12 (Wu & Nacu 2010) and only alignments with > 90% identity were kept. The resulting BAM file was reverted as a FASTA file and replaced the Trinity short-read-based transcriptome assembly in the Maker pipeline. While IsoSeq data was acquired from adult males only, short-read RNA-Seq data was generated from a pool of developmental stages and sexes (see Palmieri et al. 2015 for details), making it useful for isoform detection. We included this data in our annotation procedure as an additional file by mapping it against the v2 genome using gsnap r2017-10-12 (Wu & Nacu 2010) with default parameters. The resulting BAM file was filtered for proper pairs using SAMtools 1.5 (Li et al. 2009). Since transcript reconstruction can be confounded by high coverage (Palmieri et al. 2012), a subset of randomly sampled 50M read pairs was used in Cufflinks 2.2.1 (Trapnell et al. 2010) with default parameters. The resulting transcript file was converted from gtf to gff3 format and used as est_gff input in Maker.

To sum up, the modified annotation pipeline contained two differences compared to the initial one: (i) using IsoSeq data instead of Trinity assembly, and (ii) adding the Cufflinks transcriptome assembly from short read data as additional input file. All other parameters, options and input files (Augustus de novo gene prediction, protein sequence data) were similar between the two pipelines.

### Data availability

All PacBio data (DNA sequencing and IsoSeq) and the *D. simulans* M252 v2 genome assembly are available from the European Nucleotide Archive (ENA project ID PRJEB28741). The output of the IsoSeq pipeline (fastq files) and annotation tracks are available from Dryad.

## Results & Discussion

### DNA long read sequencing

We sequenced DNA of an inbred *D. simulans* line from Madagascar using 14 SMRTcells on a PacBio RS II. After filtering and adapter removal, this led to 8 Gb of raw data spanning 745,625 subreads with a mean read length of 6,410 bp, the longest subread reached 48,980 bp and the NR50 was 9,125 bp (NR50 is the read length such that 50% of the total sequence is contained within reads of at least this length). Mean coverage of the v1 reference genome (Palmieri et al. 2015) was ~ 50× after mapping subreads with BWA mem (Li 2013, with option -x pacbio).

### Benchmarking of assembly algorithms

We ran seven different algorithms to assemble the genome (Table 1). While all resulting assemblies were fairly similar size-wise (124 to 135 Mb), they differed in terms of fragmentation (number of contigs and N50 respectively ranging from 311 to 107,481 and from 610 kb to 5.37 Mb). Alignment of assemblies to the *D. melanogaster* reference genome r6.03 revealed all algorithms but Falcon, Canu and SPAdes produced errors with this data set. Since HGAP was developed for bacterial genome assembly, it is not optimized for eukaryote genomes. miniasm do not perform any consensus step and as such may erroneously assemble reads, and the Racon polishing step is not sufficient to uncover these assembly errors. The MaSuRCA assembly contained an assembly error on the longest contig (hence its size, Table 1) and overall the output of hybrid assemblers was much more fragmented than for PacBio-only assemblers, since many Illumina-only contigs inflated the assembly. In line with a recent benchmark study (Jayakumar & Sakakibara 2017), Falcon and Canu overall provided the best assemblies, which were both polished using long- and short-read data (Quiver and Pilon, two iterations each). Polished assemblies were merged using Quickmerge, using the Canu assembly as query. This merged assembly encompassed 127.41 Mb on 305 contigs, with a N50 of 5.38 Mb. Interestingly, the merged assembly was only marginally better in terms of contiguity and total size, while bigger improvement is usually observed with this tool (e.g., Chakraborty et al. 2016; Mahajan et al. 2018). This would be expected if both assemblies were already fairly similar, as suggested by their statistics.

**Table 1.**
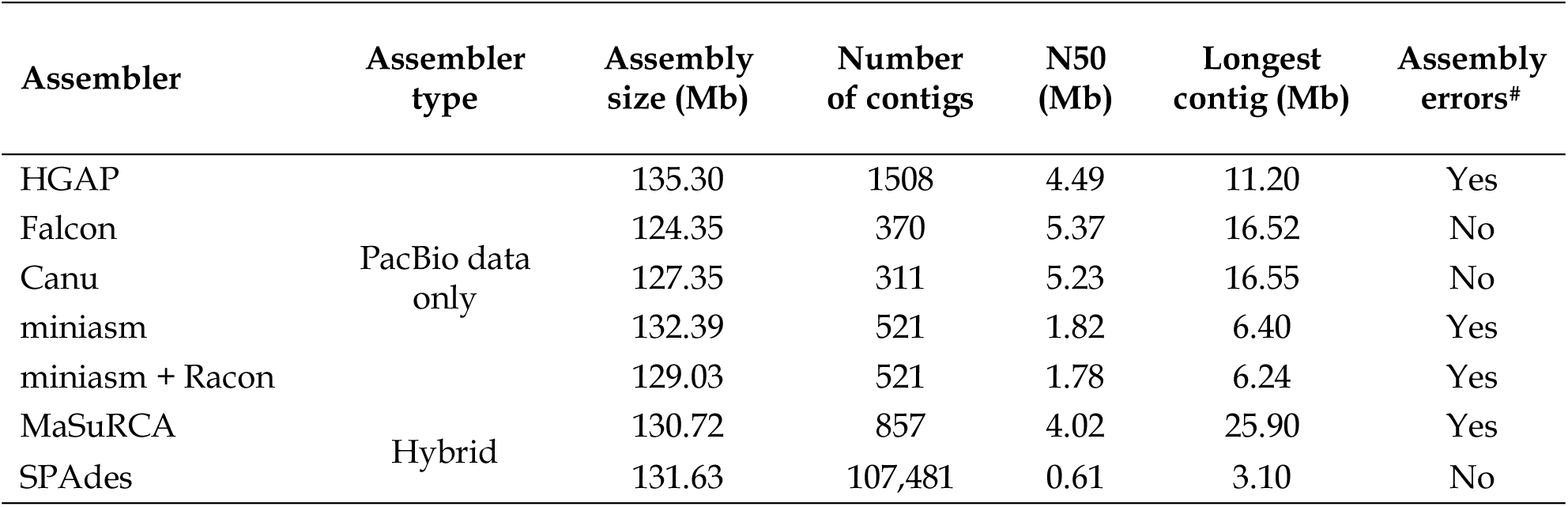
Assembly statistics obtained for *D. simulans* using seven different algorithms with ~ 50× PacBio data. Hybrid assemblers were run using this data in conjunction with > 250× Illumina paired-end data from Palmieri et al. (2015). ^#^: Assembly errors are defined as contigs aligning on two different *D. melanogaster* chromosome arms.

Small indels are the most frequent sequencing errors associated to the PacBio technology (Rhoads & Au 2015). Polishing the merged assembly with both PacBio (twice, using Quiver) and Illumina (twice, using Pilon) reads greatly reduced these errors (number of indels per 100 kb: unpolished = 136; polished = 5.03; 27-fold reduction). A significant fraction of mismatches was still present after polishing (number of mismatches per 100 kb: unpolished = 25.4; polished = 19.1) However, these values were obtained by comparing both assembly versions assuming the sequence of the v1 assembly was error-free, which is highly unlikely.

### *Anchoring on the* D. melanogaster *reference genome*

To achieve a chromosome-level assembly, polished contigs were anchored on the *D. melanogaster* r6 reference using nucmer. After alignment, the number of contigs per chromosome varied from two (chromosome 4) to 26 (X chromosome, Table 2). Compared to the v1 assembly, our PacBio-based v2 assembly contained more than six Mb of novel sequence, among which four were located on major chromosome arms. Overall our v2 assembly was 127.32 Mb, with almost five percent increase in size compared to the v1. Most of this novel sequence located within repeat-rich, telomeric or peri-centromeric regions (Figure S1), for which assembly from short-read data is challenging (van Dijk et al. 2018).

**Table 2.** Assembly size comparison of the two *D. simulans* genome versions (v1: Illumina-based, v2: PacBio-based) after anchoring of the PacBio contigs on the *D. melanogaster* reference genome r6. Δ size: size difference between v1 and v2 assemblies.

| Chromosome | Size (Mb) | | $\Delta$ size (Mb) | Number of contigs | |
| --- | --- | --- | --- | --- | --- |
|  | v1 | v2 |  | v1 | v2 |
| X | 20.62 | 21.52 | 0.90 | 412 | 26 |
| 2L | 21.09 | 21.87 | 0.78 | 491 | 8 |
| 2R | 18.98 | 19.92 | 0.94 | 307 | 16 |
| 3L | 22.25 | 22.88 | 0.63 | 229 | 10 |
| 3R | 26.97 | 27.78 | 0.81 | 265 | 15 |
| 4 | 1.10 | 1.12 | 0.02 | 42 | 2 |
| Sum | 111.01 | 115.08 | 4.07 |  |  |
| Unassembled | 10.19 | 12.23 | 2.05 | 2941 | 228 |
| Overall sum | 121.20 | 127.32 | 6.12 |  |  |

While the SMRTbell library was built using a pool of larvae (i.e., both sexes), we did not manage to assemble the Y chromosome, which is highly repetitive (Mahajan et al. 2018). The relative low amount of Y-linked sequences expected in our library compared to autosomes (hence, low coverage) could have been a major impediment to take on this task.

### Assembly quality

Nonproper read pairs could reveal the presence of misassemblies in the reference genome. We recovered the ratio of nonproper pairs in non-overlapping, 10 kb windows using an independent pool of isofemale lines collected in Florida which were aligned on both the v1 and v2 reference genomes. For all major chromosome arms, the mean ratio was smaller for the v2 assembly (see Table S1), and especially for the X chromosome (v1: 4.57%; v2: 3.59%). We plotted the ratio of nonproper pairs per window to obtain a genome-wide distribution of misassemblies for both versions (Figure 1). Most of the assembly errors found in the v1 along the X chromosome and in peri-centromeric regions were actually corrected in the v2, since these loci were assembled with less, longer contigs (Table 2). However, the v1 assembly was fairly good outside of these regions and less improvement was observed in the v2, apart from a region between 4 and 5 Mb on chromosome 2L. Since the v1 assembly was comparable to both the FlyBase r.14 (Begun et al. 2007) and the Hu et al. (Hu et al. 2013) assemblies in terms of quality (Palmieri et al. 2015), this suggests the v2 also outperforms these assemblies in terms of contiguity and assembly correctness.

**Figure 1.**
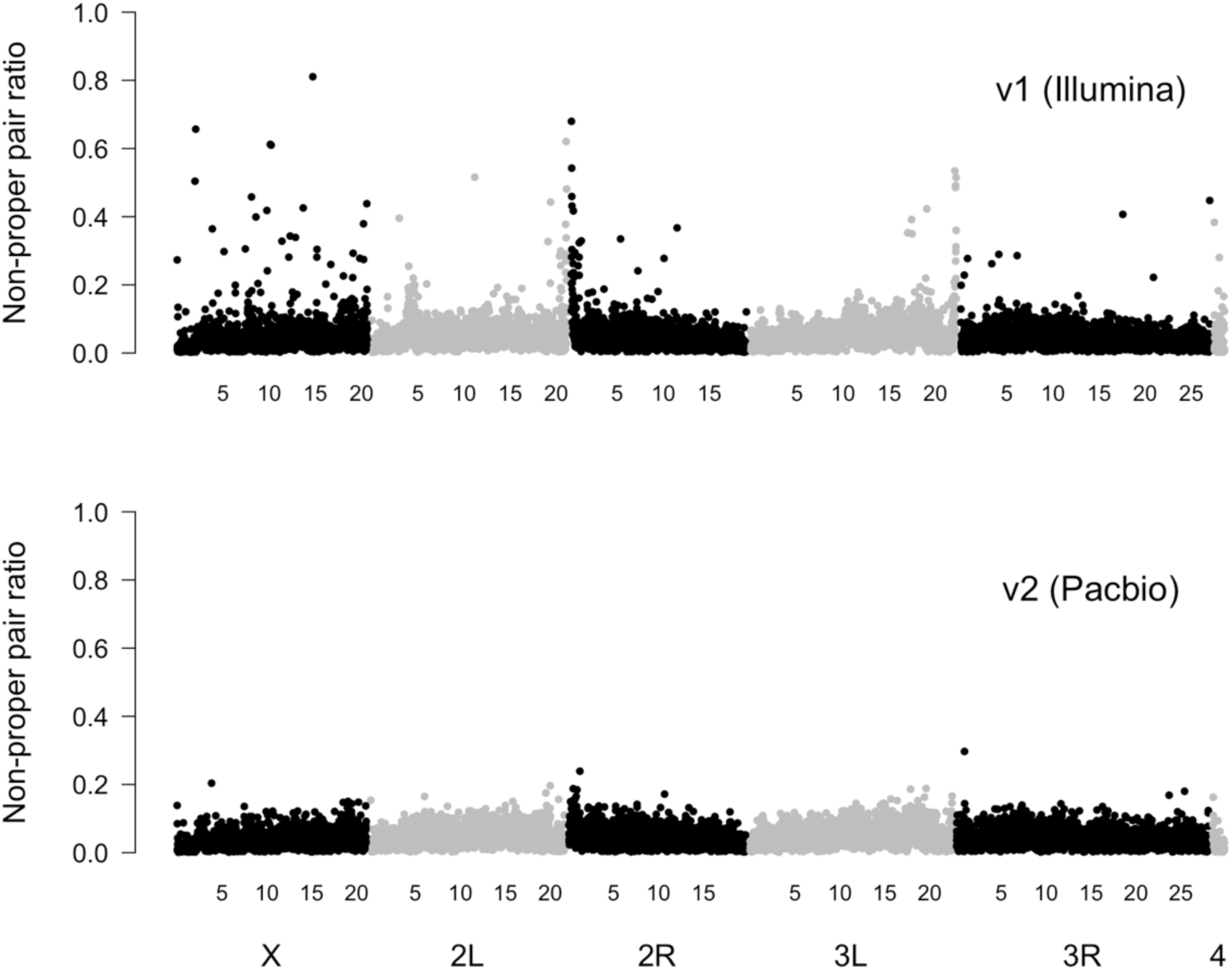
Improved contiguity of the PacBio assembly. The ratio of non-proper Illumina read pairs was recovered in non-sliding 10 kb windows along the genome for the same dataset aligned against the v1, Illumina-based assembly (Palmieri et al. 2015, upper panel) and the v2, PacBio-based assembly (lower panel). Coordinates on the x-axis are in Mb.

We assessed assembly completeness through the BUSCO analysis, using the Diptera gene set (*n* = 2799 orthologs). Our v2 assembly contained 98.7% of the orthologs among which 98.2% were complete and in single-copy, while 0.6% were detected as fragmented. Only 0.7% of the orthologs were not found in our assembly. We ran the BUSCO analysis on the *D. melanogaster* r6.17 reference and completeness of both genomes was comparable (Table 3).

**Table 3.**
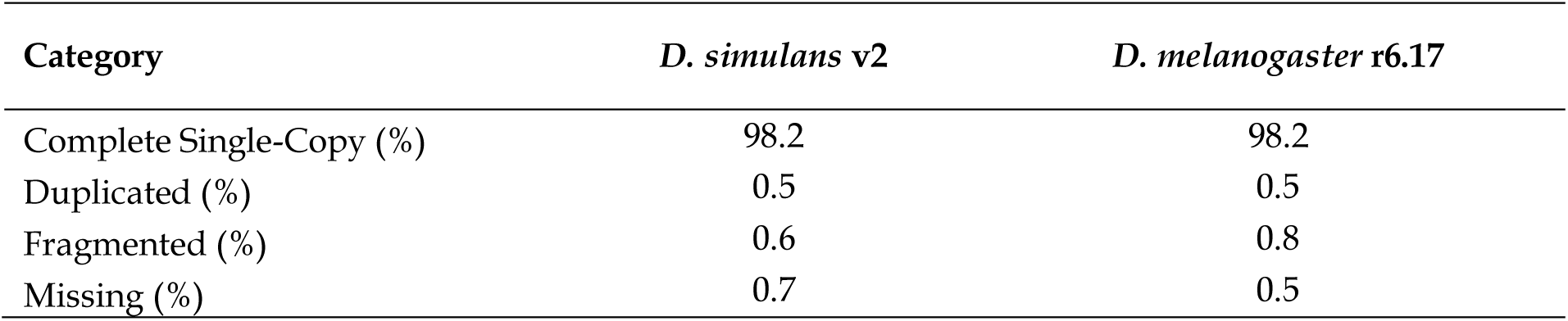
Gene content assessment for *D. simulans* v2 and *D. melanogaster* r6.17 assemblies using the BUSCO (Benchmarking Universal Single-Copy Ortholog) Diptera gene set (*n* = 2799 orthologs).

| Category | *D. simulans* v2 | *D. melanogaster* r6.17 |
| --- | --- | --- |
| Complete Single-Copy (%) | 98.2 | 98.2 |
| Duplicated (%) | 0.5 | 0.5 |
| Fragmented (%) | 0.6 | 0.8 |
| Missing (%) | 0.7 | 0.5 |

### RNA long read sequencing

We extracted RNA from two pools of *D. simulans* males from the M252 strain and sequenced two SMRT libraries using PacBio. Each cDNA library produced ~ 5 Gb of raw data and 245,000 subreads, with a mean read length of 1,916 bp. The IsoSeq pipeline was run on the two merged libraries to produce 28,250 high and 115,918 low quality isoforms with a mean length of 2,484 bp. The two sets of isoforms were pooled and mapped against the v2 reference genome and isoforms aligning with at least 90% identity were kept for the annotation step.

### Annotation

We used the Maker pipeline to annotate the genome by combining ab initio predictions from Augustus, RNAseq data from Palmieri et al. (2015) and protein sequence data from *D. melanogaster* and *D. simulans.* Visual comparison with the *D. melanogaster* r6.17 annotation in FlyBase revealed two issues: nested gene structures were misidentified (Figure 2), and gene fusion events occurred when UTRs overlapped. Following Maker’s guidelines (see Methods) did not solve these issues. Inclusion of the IsoSeq data in the annotation pipeline allowed us to reconstruct correctly such gene structures while limiting gene fusion events (Figure 2). This improved pipeline recovered 13,422 genes and 18,301 transcripts (mean number of isoforms per gene: 1.36). The mean gene length was 5,335 bp and the mean number of exons per gene was 2.98, with a mean exon length of 492 bp and a mean intron length of 967 bp. Overall, genes accounted for 56.4% of the genome. These numbers are in line with previous observations made for *D. simulans* (Begun et al. 2007; Hu et al. 2013; Palmieri et al. 2015).

**Figure 2.**
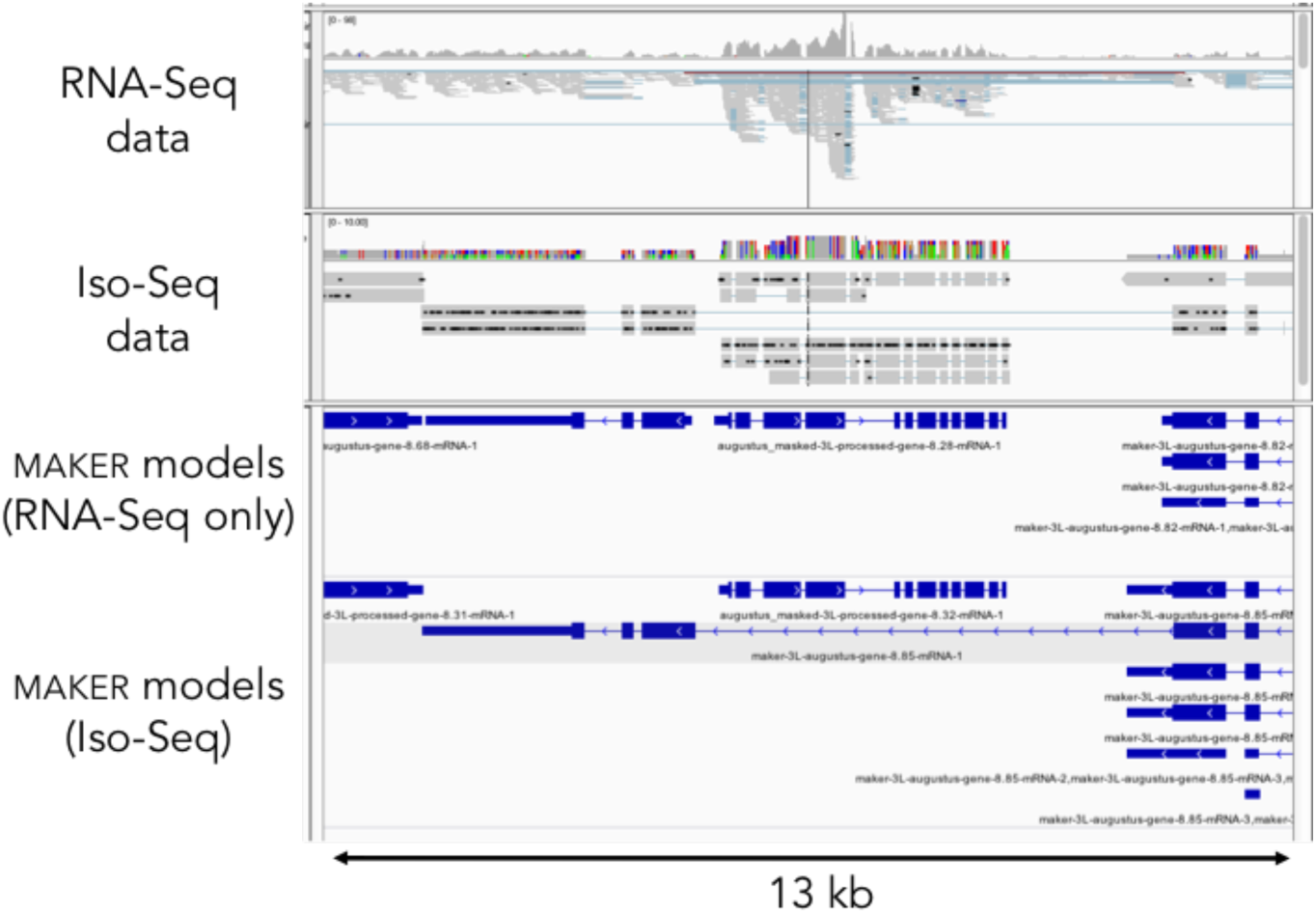
IsoSeq data helps automated genome annotation. Maker models (third panel from top, in IGV) built from RNA-Seq data (top panel) cannot resolve the nested gene structure, while Maker models (bottom panel) built from IsoSeq data (second panel from top) can.

Maker uses the Annotation Edit Distance (AED) to measure the agreement between ab initio predictions and empirical evidence for each gene model (0 indicates perfect agreement whereas 1 indicates no support of empirical data). Our gene models had a mean AED of 0.18 (median: 0.13) and 98% of them had a distance smaller than 0.5. Finally, we could validate 90% of our gene models by best reciprocal BLAST against *D. melanogaster* r6.17 transcripts. These results suggest most of our annotations are of high quality. Our study demonstrates that PacBio sequencing of full-length transcripts associated with the IsoSeq pipeline represent a sound approach to build high-quality annotations. Depending on the size of the transcriptome, and with the decreasing costs of PacBio sequencing, this could represent a promising strategy also in non-model organisms.

## Conclusion

This work provides the first complete long-read-based assembly of a *D. simulans* genome and adds to the platinium-grade assemblies available for *Drosophila* (Chakraborty et al. 2018; Mahajan et al. 2018; Miller et al. 2018). With a high contiguity level and a gene content completeness comparable to the *D. melanogaster* genome, this will provide a useful resource for genomic studies. The better assembly of peri-centromeric and telomeric regions could also allow to investigate the evolution of transposable elements at an unprecedented scale. Finally, including IsoSeq data in the annotation improved characterization of complex gene structures and should thus provide a better resolution for future RNAseq studies in *D. simulans*.

## Acknowledgments

I wish to acknowledge Viola Nolte for library preparation, Lukas Endler for assisting with SMRTanalysis installation, Heinz Ekker from the VBCF for running the IsoSeq pipeline and Christian Schlöterrer and colleagues from the Institut für Populationsgenetik for constructive feedback. This research was supported by the European Research Council grant “Archadapt” awarded to Christian Schlöterrer and a Bright Spark grant of the vetmeduni Vienna awarded to PN. The funders had no role in study design, data collection and analysis, decision to publish, or preparation of the manuscript. I declare no conflict of interest.

## Supplementary data

**Table S1.**
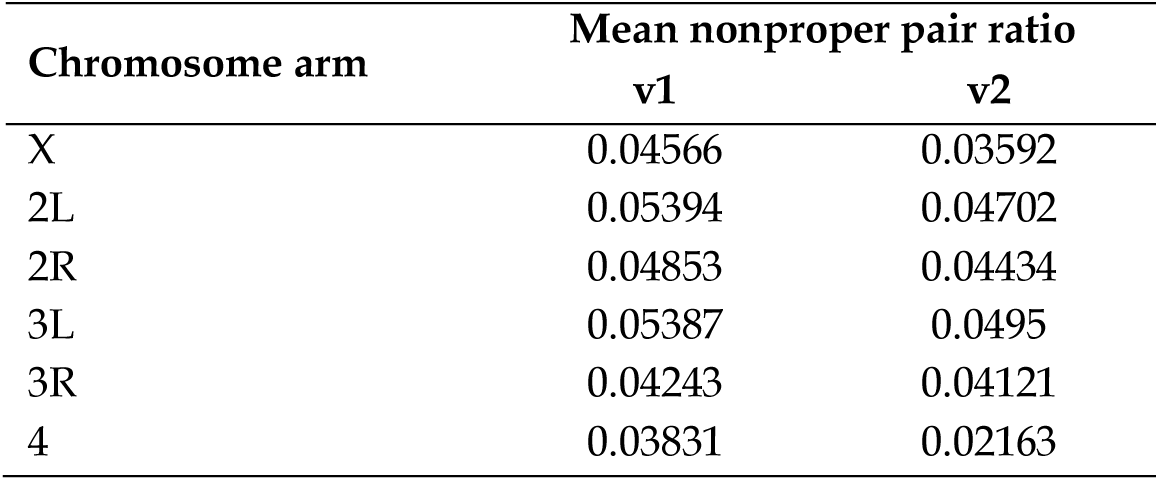
Mean nonproper pair ratio for the different major chromosome arms obtained from the same data set after mapping to both reference genomes (v1: Illumina-based; v2: PacBio-based).

**Figure S1.**
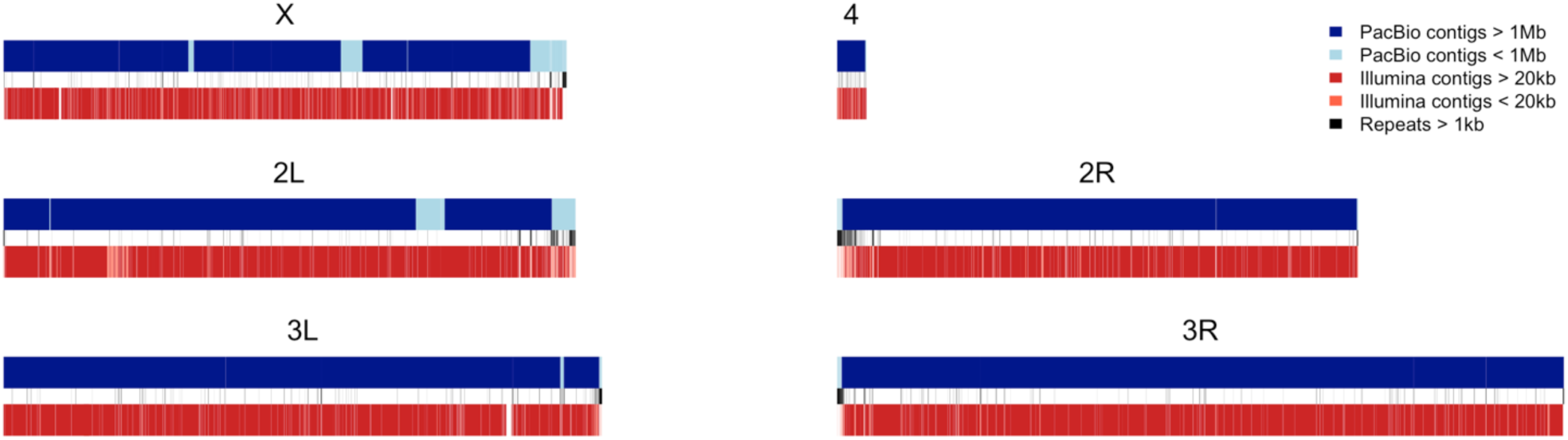
Mapping of PacBio (blue) and Illumina (red) contigs on major chromosomes of the v2 reference genome. Repeats (> 1kb) were identified by RepeatMasker and are indicated in black. Illumina contigs are from Palmieri et al. (2015).

### File S1

Optimized config file used for the Falcon assembler.

~~~
[General]
job_type = local

input_fofn = input.fofn

input_type = raw

length_cutoff = –1

genome_size = 162000000
seed_coverage = 30

length_cutoff_pr = 2000

target = assembly

sge_option_da = –pe smp 8 –q 32
sge_option_la = –pe smp 8 –q 32
sge_option_pda = –pe smp 8 –q 32
sge_option_pla = –pe smp 8 –q 32
sge_option_fc = –pe smp 8 –q 32
sge_option_cns = –pe smp 8 –q 32

default_concurrent_jobs = 8
pa_concurrent_jobs = 8
cns_concurrent_jobs = 8
ovlp_concurrent_jobs = 8

# preassembly
pa_DBsplit_option = –a –×500 –s100
pa_HPCdaligner_option = –v –B128 –t16 –M32 –e.70 –l1000 –s100 –k18 –h480 –w8

# error correction
falcon_sense_option = ‐‐output_multi ‐‐min_idt 0.70 ‐‐min_cov 4 ‐‐max_n_read 200 ‐‐
n_core 8
falcon_sense_skip_contained = True

# overlapping of corrected reads
ovlp_DBsplit_option = –a –×500 –s100
ovlp_HPCdaligner_option = –v –B128 –M32 –h1024 –e.96 –l1000 –s100 –k24

overlap_filtering_setting = ‐‐max_diff 40 ‐‐max_cov 45 ‐‐min_cov 1 ‐‐n_core 12
~~~

